# Metabolic crosstalk between hydroxylated monoterpenes and salicylic acid in tomato defence response against *Pseudomonas syringae* pv *tomato*

**DOI:** 10.1101/2023.05.05.539605

**Authors:** J Pérez-Pérez, S Minguillón, E Kabbas-Piñango, C Payá, L Campos, M Rodríguez-Concepción, I Rodrigo, JM Bellés, MP López-Gresa, P Lisón

## Abstract

Hydroxylated monoterpenes (HMTPs) are differentially emitted by tomato plants efficiently resisting a bacterial infection. We have studied the defensive role of these volatiles in the tomato response to bacteria, whose main entrance are stomata apertures. Treatments with some HMTPs resulted in stomatal closure and *PR1* induction. Particularly, α-terpineol induced stomatal closure in a salicylic (SA) and abscisic acid-independent manner, and conferred resistance to bacteria. Interestingly, transgenic tomato plants overexpressing or silencing the monoterpene synthase *MTS1,* which displayed alterations in the emission of HMTPs, exhibited changes in the stomatal aperture but not in plant resistance. Measures of both 2-C-methyl-D-erythritol-2,4-cyclopyrophosphate (MEcPP) and SA levels, revealed a competition for MEcPP by the methylerythritol phosphate (MEP) pathway and the SA biosynthesis activation, thus explaining the absence of phenotype in transgenic plants. These results were confirmed by chemical inhibition or activation of the MEP pathway. Besides, treatments with BTH, a SA functional analogue, conferred enhanced resistance in transgenic tomato plants overexpressing *MTS1.* Finally, plants overexpressing *MTS1* induced *PR1* and stomata closure in neighbouring plants. Our results confirm the role of HMTPs in both intra and inter-plant immune signalling, and reveal a metabolic crosstalk between the MEP and SA pathways in tomato plants.

**Graphical Abstract:** Metabolic crosstalk between hydroxylated monoterpenes and salicylic acid in tomato defence response against Pseudomonas syringae pv tomato.
Created with BioRender.com.

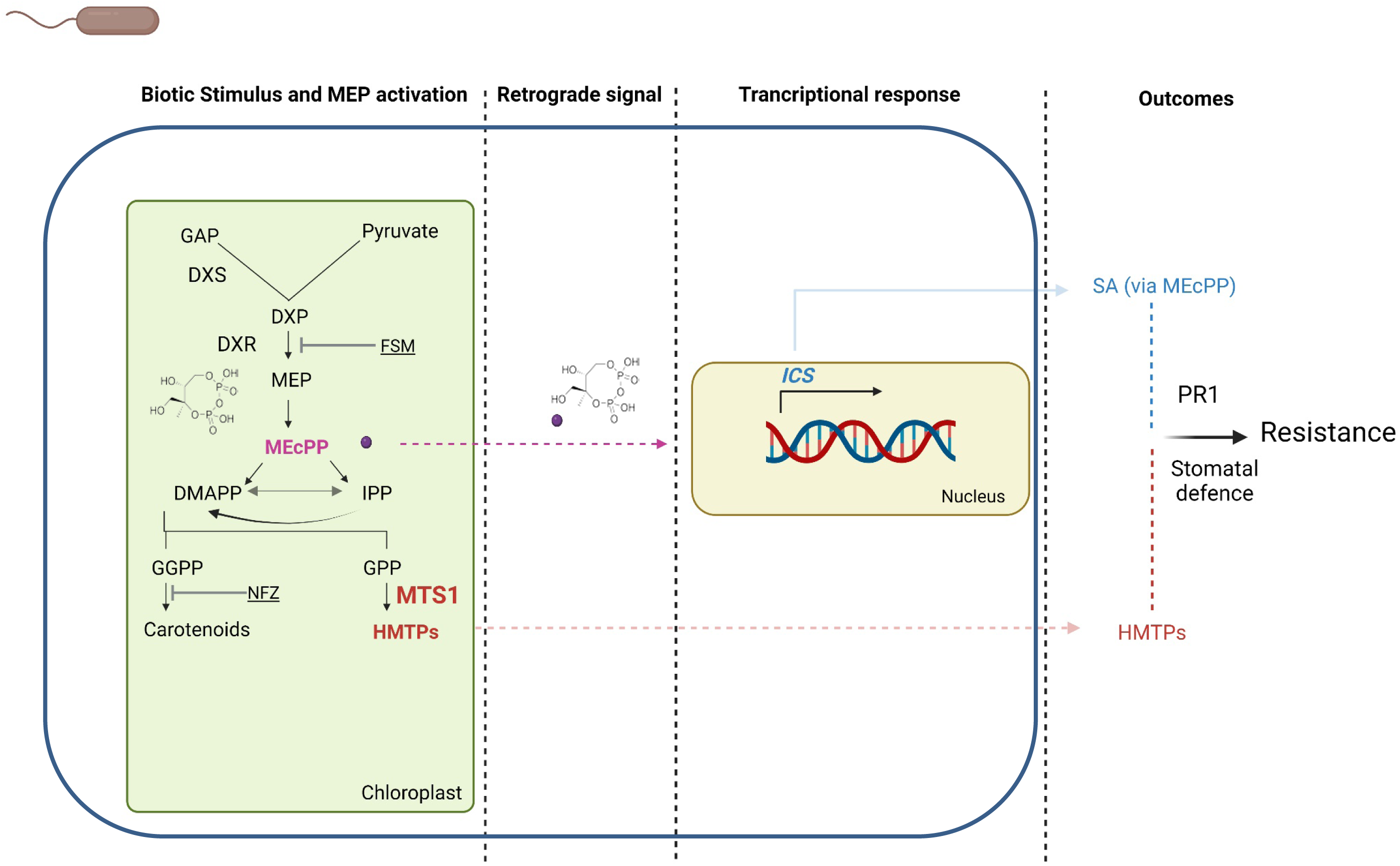

## INTRODUCTION

Understanding the defensive signalling pathways in plants has allowed the discovery of new resistance-inducing compounds for the agrochemical sector. These compounds may act directly as powerful antioxidants, antibacterial or antifungal agents against the pathogen, or act indirectly by activating the plant defence response. In addition to alkaloids and phenolic compounds, some volatile organic compounds (VOCs) belong to this group of defensive molecules.

One of the most diverse types of VOCs are terpenoids, also known as isoprenoids, and include a very extensive and diverse set of molecules formed from repeating units of 5-carbon (C5) building blocks, named isopentenyl diphosphate and dimethylallyl diphosphate, which are obtained from mevalonic acid (MVA) or methylerythritol phosphate (MEP) in the cytosol or plastids, respectively. Particularly, monoterpenes consist of two building blocks (C10) and can be modified with different functional groups to form monoterpenoids, which include the hydroxylated monoterpenes (HMTPs). TPS catalyze the synthesis of isoprenoids, being responsible for the diversity of terpenoids found in Nature (Tholl and Lee, 2011). The modification of monoterpenes to produce HMTPs could be carried out through the cytochrome P450 monooxygenases (Karunanithi and Zerbe; 2019). The analysis of the updated tomato genome (2017 version of v. SL3.0) has revealed that there are 34 full-length *TPS* genes and 18 pseudo *TPS* genes. The biochemical analysis has identified the catalytic activities of all the enzymes encoded by all 34 *TPS* genes: an isoprene (C5) synthase, 10 exclusively or predominantly monoterpene (C10) synthases, 17 sesquiterpene (C15) synthases, and 6 diterpene (C20) synthases (Zhouand Pichersky, 2020). Among TPS, the recombinant protein of the monoterpene synthase MTS1 (Graphical Abstract) produces the monoterpenoid β-linalool but also the sesquiterpenoid β-nerolidol, generating its overexpression increased levels of linalool in tomato plants. This gene is induced by insects, wounding and jasmonic acid (JA)-treatment (Van Schie et al., 2007). Therefore, the defensive role of monoterpenes has been classically associated with plant-herbivore interaction, although the interest on its role in plant defence against pathogens is increasing (Vlot et al., 2021).

A non-targeted metabolomic analysis was performed using gas chromatography coupled to mass spectrometry to identify VOCs differentially emitted by Rio Grande (RG) tomato plants carrying the *Pto* resistance gene, infected by an avirulent strain of *Pseudomonas syringæ* pv. *tomato*. The analysis of the specific volatilome from these plants, displaying the so-called Effector-Triggered Immunity (ETI; Jones and Dangl, 2006), revealed that the aroma of resistance was mainly formed by (Z)-3-hexenol esters -including (Z)-3-hexenyl butanoate (HB)-as well as some HMTPs, such as α-terpineol, 4-terpineol and linalool (López-Gresa et al., 2017). The defensive role of HB has been already demonstrated, since it produces stomatal closure and induce the defensive response, thus preventing the entry of bacteria (López-Gresa et al., 2018). This compound was patented (Lisón et al., 2018) because of its potential uses in agriculture against biotic and abiotic stresses (Payá et al. 2020; Payá et al., 2023 under review). However, the role of HMTPs in tomato immunity remains unknown. HMTPs, and particularly linalool, are synthesized through the MEP plastid pathway by the action of *MTS1*, which has been described to be induced during ETI in tomato plants (López-Gresa et al., 2017).

The defence response upon avirulent *Pseudomonas syringæ* pv. *tomato* infection also includes the activation of salicylic acid (SA) biosynthesis (Robert-Seilaniantz et al., 2011). This phytohormone is involved in different physiological and biochemical processes and it is well characterized as a signalling molecule for the induction of different pathways that lead to an enhance in plant resistance (Klessig et al., 2018). However, the mechanisms of SA signal transduction pathways have not been elucidated yet (Pokotylo, et al 2019). This phenolic compound is biosynthesized in plants from phenylalanine through the route of the phenylpropanoids (PAL pathway) or from isochorismate (IC pathway). Loss-of-function of some genes from both pathways results in an increased plant susceptibility to pathogens. Nevertheless, isochorismate synthase (ICS1) is the main enzyme of SA biosynthesis in biotic responses and produces 90% of its levels under biotic stress (Wildermuth et al., 2001; Zhang and Li, 2019). In plants, IC is conjugated to the amino acid L-glutamate by an isochorismoyl-9-glutamate (IC-9-Glu) that can spontaneously break down into SA. Besides, EPS1 (Enhanced *Pseudomonas* Susceptibility 1), an IC-9-Glu pyruvoyl-glutamate lyase, can enhance this process more effectively (Zeier, 2021). To avoid its toxic effects caused by its accumulation, SA can be chemically modified into different derivatives, through glycosylation, methylation, sulfonation, amino acid conjugation, and hydroxylation. Particularly, most of the SA present in the plant is glycosylated into SA 2-O-β-D-glucoside (SAG). In addition, SA can be methylated to form the volatile methyl salicylate (MeSA) or hydroxylated to form gentisic acid (GA) by the action of S5H (Salicylic acid 5-Hydroxylase; Ding and Ding 2020; Payá et al., 2022).

The differential emission of HMTPs in an avirulent bacterial infection, as well as the observed induction of the monoterpene synthase *MTS1* (López-Gresa et al., 2017) prompted us to delve into the possible defensive role of HMTPs in tomato plants. In this context, the general objective of this work is to study the defensive role and the mode of action of HMTPs, including the possible interrelation with SA, in the tomato-bacteria interaction.

## RESULTS

### HMTPs Activate the Plant Defence Response

Stomata play a critical role in restricting bacterial invasion as part of the plant immune system (Underwood, 2007). In addition to biotic stress, stomata also respond to several volatile compounds, some of them with defensive activity (Lopez-Gresa et al. 2018). In order to explore the defensive role of HMTPs whose differentially emission is triggered by ETI, MoneyMaker (MM) tomato plants were treated either with 5 µM α-terpineol, 4-terpineol or linalool, and both stomata closure and expression of *PR1*, the main marker gene of SA-mediated plant response to biotrophic attack (Rodrigo et al., 1993; Tornero et al., 1993), were analysed. Besides, treatments with the non-hydroxylated monoterpene limonene were also performed. As observed in Figure 1A, a significant stomatal closure occurred in α-terpineol, 4-terpineol and linalool but not in limonene-treated plants. In a similar manner, HMTPs treatments also produced a significant induction of *PR1* expression, whilst limonene had no clear effect (Figure 1B). Our results appear to indicate that both stomatal closure and activation of *PR1* are specifically triggered by the hydroxylated forms of monoterpenes, being α-terpineol selected for further studies, since it was the compound producing the higher stomatal closure.

**Figure 1.**
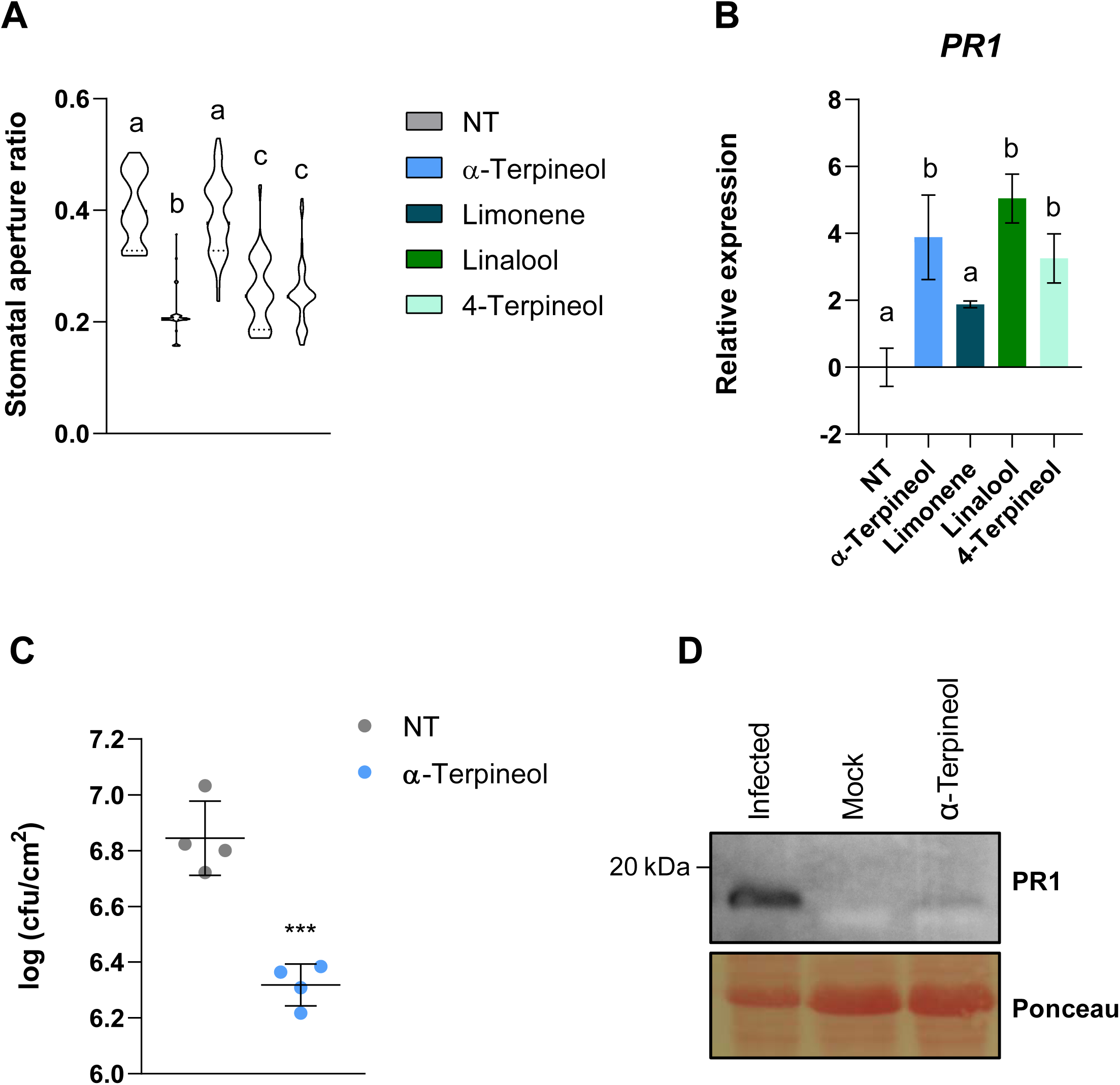
Effect of monoterpenoid treatments on the defensive response of MoneyMaker tomato plants. **A)** Stomatal aperture ratio of non treated (NT) tomato plants or treated with α-terpineol, limonene, linalool and 4-terpineol. Violin plots represent the stomatal aperture ratio for each treatment of a total of 40 stomata from three biological replicates. Different letters indicate statistically significant differences for each treatment (*p* < 0.05). **B)** Real-time qPCR analysis of the tomato *PR1* gene expression in non treated (NT) tomato plants or treated with α-terpineol, limonene, linalool and 4-terpineol. Values were normalized to Actin gene. Expression levels are represented as mean ± SD of three biological replicates of one representative experiment. Letters represent statistically significant differences (*p* < 0.05) between treatments. **C)** Bacterial content in leaves. Tomato plants were pre-treated with 5 µM α-terpineol or non-treated (NT), and then inoculated by immersion one day later. Bacterial growth measurements were performed 24 hours after bacterial infection. Data are presented as mean ± SD of a representative experiment (n=4). Statistically significant differences (*p* <0.05) between treated and non-treated plants are represented by different letters. **D)** Western blot analysis of PR1 (14 kDa) accumulation in tomato leaves after 24 h of α-terpineol treatments.

To confirm the defensive role of α-terpineol, MM tomato plants were pre-treated with 5 µM α-terpineol and subjected to *Pseudomonas syringæ* pv. *tomato* DC3000 (*Pst*) infection by immersion one day later (see materials and methods). A significant induction of resistance was observed in α-terpineol treated tomato plants after 24h of bacterial inoculation, when compared with the non-treated plants (Figure 1C). The efficacy of the pre-treatment was also tested in the experiment, confirming the activation of PR1 by *Western blot* analyses (Figure 1D).

### α-Terpineol Induces Stomatal Closure in a SA/ABA-Independent Manner

To better explore the function of α-terpineol in stomatal closure, weight loss was measured in tomato seedlings after water or α-terpineol treatments during 120 min (see Methods). As shown in Figure 2A, α-terpineol-treated plants statistically retained more water than non-treated plants, indicating an effective stomatal closure after the chemical treatment. We also compare the effect of α-terpineol with that caused by abscisic acid (ABA), the main phytohormone involved in stomata closure. We observed that treatments with α-terpineol, at a comparable range of concentrations as that used for ABA, can close stomata, although the observed effect was lower (Figure 2B).

**Figure 2.**
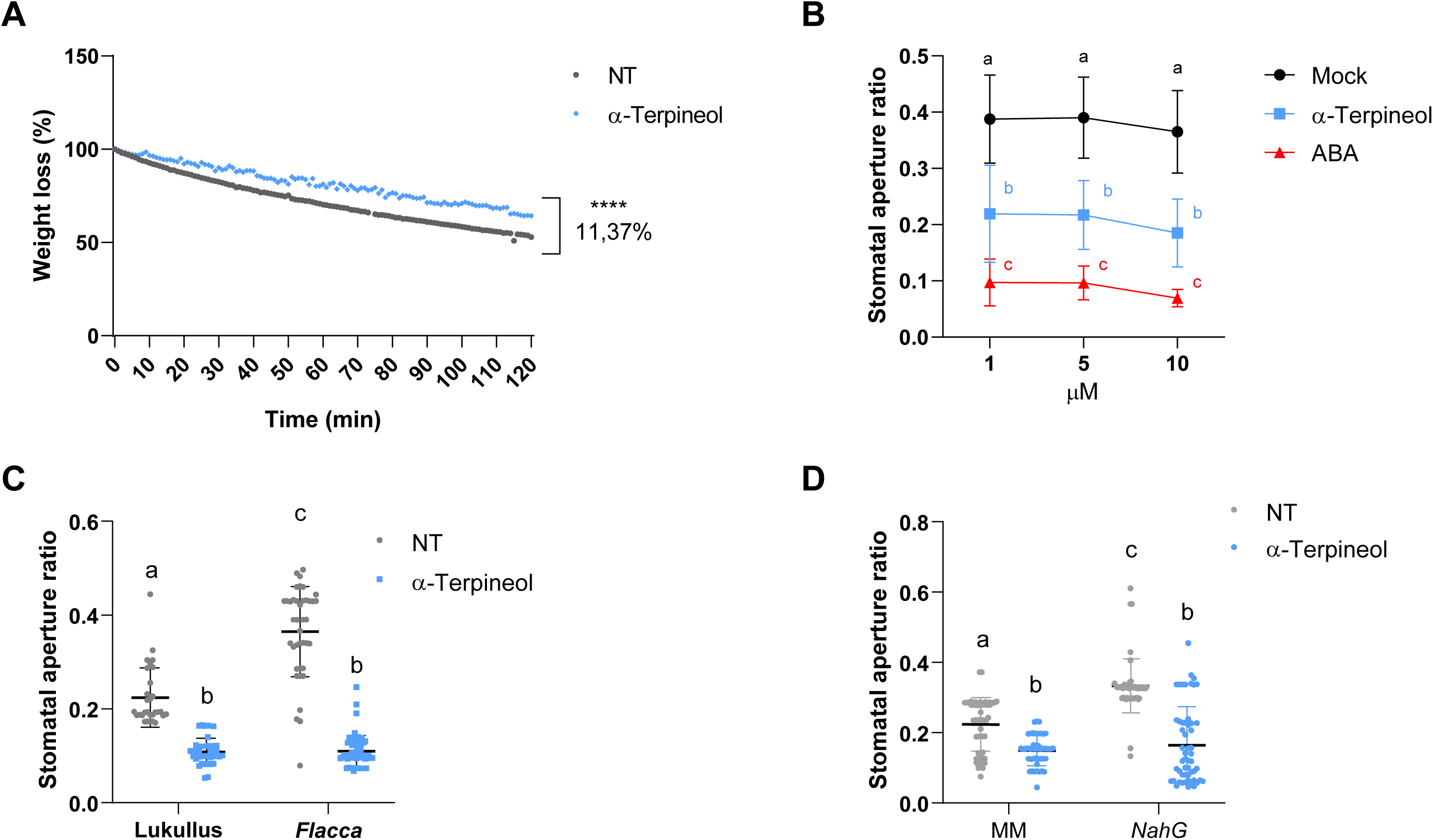
Effect of α-terpineol treatments on stomatal closure of tomato leaves. Upper panels. **A)** Weight loss of non-treated (NT) or α-terpineol treated tomato seedlings at different time points. **B)** Dose-response analysis of α-terpineol and ABA in stomatal aperture. Data represent the mean ± SD of a representative experiment (n=40). Letters indicate statistically significant differences for each treatment at each time point (*p* < 0.05). **Lower panels.** Stomatal aperture ratio mean values ± SD of three biological replicates of non-treated (NT) and α-terpineol treated tomato **C)** ABA deficient mutants (*flacca*) and the corresponding parental (*Lukullus*) and **D)** transgenic plants impaired in salicylic acid accumulation (*NahG*) and the corresponding parental MoneyMarker (MM). Letters indicate statistically significant differences from control treatments and parental plants (*p* < 0.05).

In addition to ABA, SA induces stomatal closure as a signalling molecule for plant defence responses to bacterial pathogens (Panchal and Melotto, 2017). To test if the observed terpineol-induced stomatal closure is SA or ABA dependent, we measured the effect in tomato SA-deficient *NahG* transgenic plants (Brading et al., 2000) and ABA-deficient *flacca* mutants (Bowman et al., 1984). Terpineol treatments resulted in a significant stomatal closure in *NahG* and *flacca* (Figures 2C and D), thus indicating that the effect of α-terpineol is SA and ABA-independent.

### Alteration of *MTS1* Gene Expression Levels Affects Stomatal Closure but not *Pst* Resistance

To provide genetic evidence confirming the observed defence-related effects triggered by HMTPs, tomato transgenic plants with altered levels of these volatiles were studied. Transgenic tomato plants overexpressing *MTS1* were previously described (van Schie et al., 2007) and silenced *MTS1* tomato plants were generated by following an RNAi strategy (see materials and methods; Figure 3A). Generated tomato lines *RNAi_MTS1* were characterized, and homozygous lines *RNAi_MTS1 2.1* and *RNAi_MTS1 5.1* both carrying one copy of the transgene, were selected for further studies. To characterize the response of *RNAi_MTS1* tomato plants to bacterial infection, the *MTS1* expression levels in mock-inoculated and *Pst*-infected plants were analyzed by qRT-PCR (Figure 3B). As expected, a statistically significant reduction of *MTS1* transcript was measured in *Pst*-infected *RNAi_MTS1* leaves compared to corresponding infected wild type RG.

**Figure 3.**
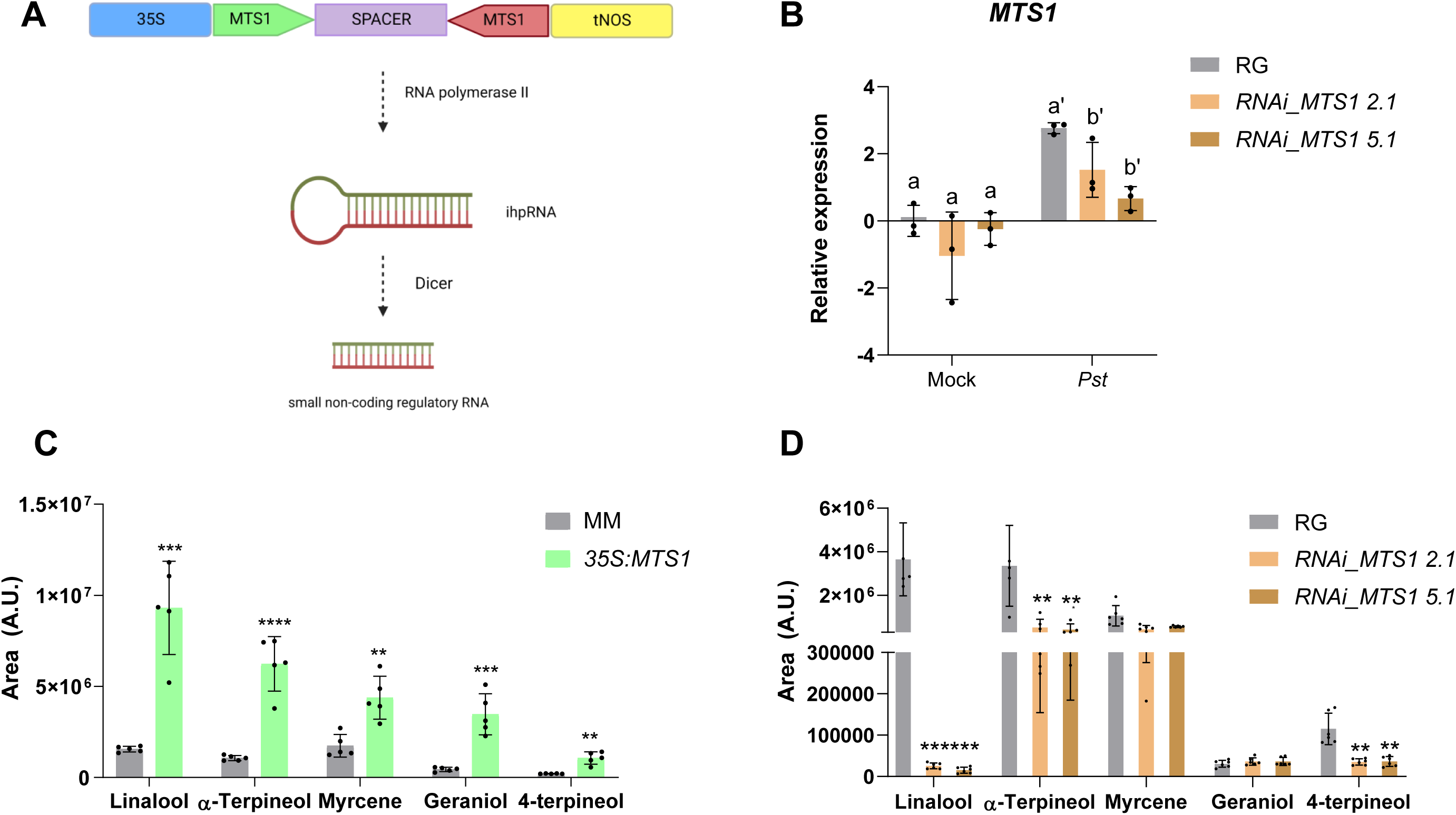
Characterization of transgenic plants with altered levels of monoterpenoids after bacterial infection. **A)** DNA construction for the generation of transgenic plants *RNAi_MTS1.* **B)** Analysis of *MTS1* expression by qRT-PCR of the different *RNAi_MTS1* transgenic tomato lines (2.1 and 5.1) and its parental (RG; Rio Grande) infected with bacterial (*Pst*) or non-inoculated (Mock). RG and transgenic plants were subjected to infection with *Pst* by immersion. Samples were taken 24 h after the bacterial infection. The qRT-PCR values were normalized with the level of expression of Actin gene. The expression levels correspond to the mean ± SD of a representative experiment (n=3). Statistically significant differences (*p* < 0.05) between genotypes and *Pst*-infected or Mock plants are represented by different letters. Targeted GC-MS for monoterpenoids was performed in tomato *35S:MTS1* leaves and their control transgenic plants with empty vector (MM; **C**), and lines of *RNAi_MTS1* 2.1 and 5.1 and their parental (RG; **D**) upon *Pst*-infection. Data are presented as means ± SD of a representative experiment (n=5). Statistically significant differences are represented with asterisk (*), double asterisk (**) and triple asterisk (***), and indicate significance differences with respect to genetic background (MM or RG) with *p* < 0.05, *p* < 0.01 and *p* < 0.001, respectively.

To determine the emission of VOCs upon a bacterial infection in the *35S:MTS1* and *RNAi_MTS1* transgenic plants, a monoterpenoids targeted analysis was performed. Figure 3C shows that the emission of monoterpenoid-type VOCs was statistically higher in the *Pst*-infected *35S:MTS1* transgenic plants compared to MM wild type plants. These results confirm the function described for *MTS1* as a monoterpene synthase and correlate with those previously described in which the basal levels of linalool reported in transgenic plants were higher than those in plants transformed with the empty vector (Van Schie et al., 2007). The chemical composition of *RNAi_MTS1* (Figure 3D) also confirms that the silencing of *MTS1* upon a *Pst* infection causes a significantly reduction of the HMTPs emission, specifically, linalool, terpinen-4-ol, and α-terpineol. Therefore, *MTS1* overxpression or silencing in infected tomato plants produced an opposite HMTPs emission.

To confirm the mode of action of HMTPs in the tomato defensive response we checked the possible correlation between HMTPs emission and stomatal closure. As shown in Figure 4A, *35S:MTS1* transgenic plants displayed a lower ratio of stomatal aperture than the corresponding control plants. This result agrees with the stomatal closure observed with exogenous α-terpineol treatments (Figure 1A), reaffirming the role of HMTPs in the regulation of stomatal closure. Also consistently, both transgenic lines silencing *MTS1*, with lower levels of HMTPs, displayed a higher aperture ratio than the wild type RG plants (Figure 4D). Thus, our results appear to indicate HMTPs can cause stomatal closure when provided exogenously but also when produced endogenously, probably being involved in stomatal immunity.

**Figure 4.**
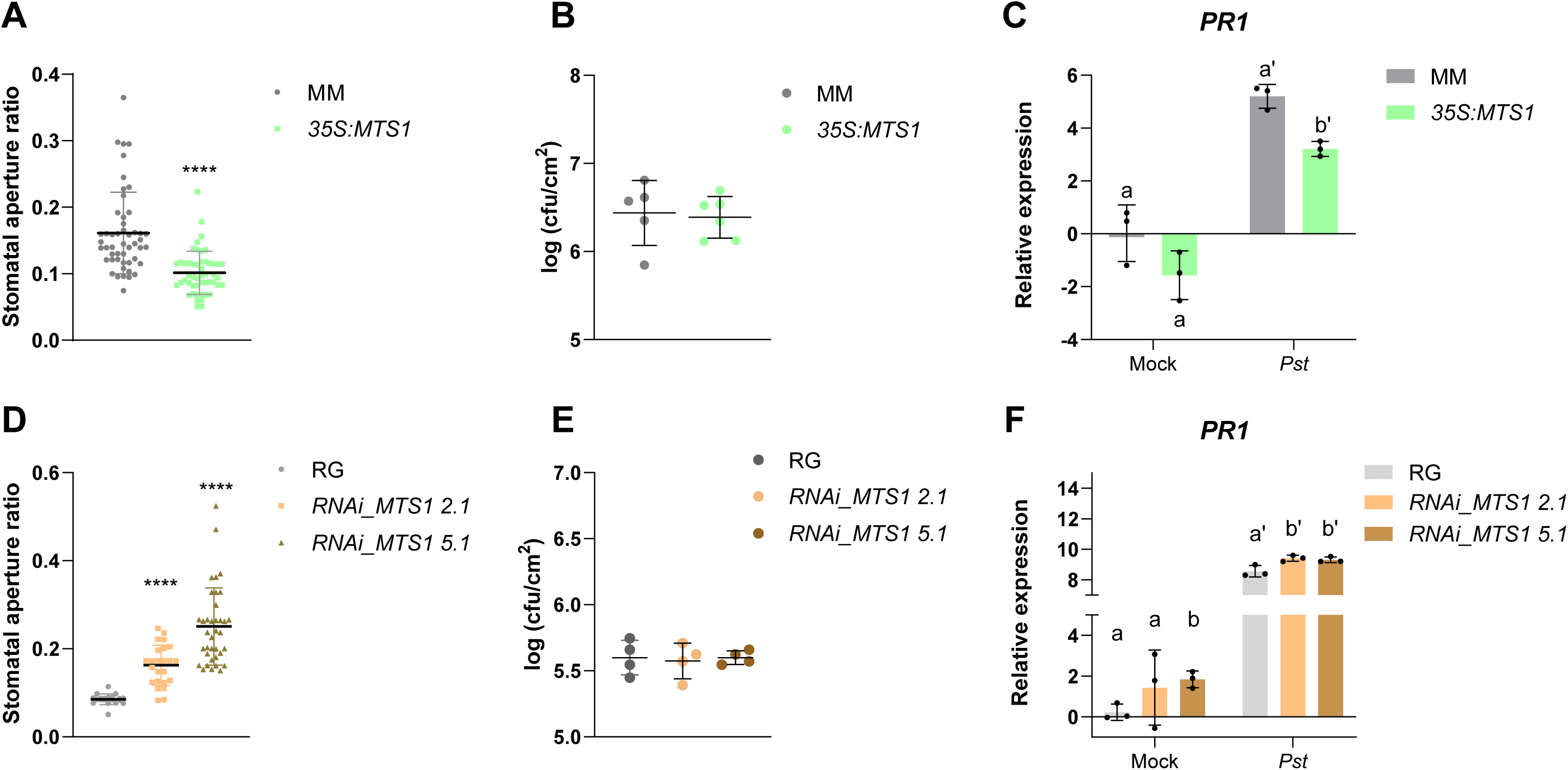
Activation of the defensive response in tomato plants with altered levels of monoterpenoids upon bacterial infection. Stomatal aperture, bacterial infectivity and *PR1* gene expression were studied for transgenic tomato lines overexpressing (upper panels, **A** to **C**) and silencing *MTS1* gene (lower panels, **D** to **F**). **Leftmost panels:** Stomatal aperture ratio mean values ± SD of a total of 40 stomata from three biological replicates are shown in **A)** for *35S:MTS1* leaves and their control transgenic plants with empty vector (MM), and in **D)** for lines of *RNAi_MTS1* 2.1 and 5.1 and their parental (RG). Asteriks (***) indicate statistically significant differences between genotypes (*p* < 0.001). **Central panels:** Growth of *Pst* are shown in leaves of **B)** *35S:MTS1* plants and their parental (MM) and **E)** both silencing lines of *RNAi_MTS1,* and their parental RG. Tomato plants were inoculated with bacterial *Pst* by immersion and leaf samples were taken 24 hours after bacterial infection. Data are presented as means ± SD of a representative experiment (n=5 and n=4, respectively). **Rightmost panels:** qRT-PCR expression analysis of the tomato *PR1* gene are shown in **C)** *35S:MTS1* plants and their parental (MM) and **F)** both silencing lines of *RNAi_MTS1* and their parental RG. Mock represents the non-inoculated plants. Values were normalized to Actin gene. Expression levels are represented as mean ± SD of three biological replicates of one representative experiment. Statistically significant differences (*p* <0.05) between genotypes and infected or mock-treated plants are represented by different letters.

To test this possibility, a bacterial infection was carried out in both transgenic plants *35S:MTS1* and *RNAi_MTS1* as in their corresponding MM and RG parentals. Surprisingly, as seen in Figure 4B, the *MTS1* overexpression line did not show enhanced resistance to *Pst,* despite having a lower stomata aperture ratio than their corresponding MM (Figure 4A). Besides, *RNAi_MTS1* tomato plants did not display a higher susceptibility (Figure 4E) despite the previous observed higher stomata aperture ratio (Figure 4D). These results are also in contrast with those obtained after exogenous treatments (Figure 1C).

Next, the expression levels of *PR1* in both infected transgenic plants were measured. As shown in Figure 4C, a statistically lower expression of *PR1* after *Pst* infection was detected in *35S:MTS1* transgenic plants when compared to that observed in MM wild type plants, which is in contrast with the previously observed HMTPs-mediated *PR1* induction (Figure 1B). Since the induction of this defence marker gene is SA dependent, these results suggested that SA signalling could be repressed under stress conditions in tomato transgenic *35S:MTS1* plants. Contrary, both *RNAi_MTS1* lines showed statistical higher levels of *PR1* expression upon *Pst* infection (Figure 4F). These results indicate that the alteration of HMTPs levels could affect the SA signalling.

### Metabolic Crosstalk Between MEP and SA Pathways in Infected Tomato Plants

HMTPs are produced from precursors supplied by the plastidial MEP pathway (Graphical Abstract). Interestingly, an intermediate of the MEP pathway, methylerythritol cyclodiphosphate (MEcPP), has been shown to transcriptionally activate *ICS*, which encodes the key enzyme of SA biosynthesis (Gil et al., 2005; Xiao et al., 2012). To study the possible competition for MEcPP to activate SA-mediated response or HMTPs biosynthesis, measures of MEcPP levels were performed in *35S:MTS1* tomato plants upon *Pst* infection (Figure 5), observing a statistically reduction of this compound in *35S:MTS1* tomato plants. This reduction in MEcPP levels, was accompanied by a significant reduction in the *ICS* expression in *35S:MTS1* infected tomato plants, when compared with the corresponding parental plants (Figure 6A). The decrease in MEcPP levels and *ICS* expression produced by the over-production of HMTPs (Figure 3C), could be the responsible for the observed lower levels of *PR1* (Figure 4C) and the absence of resistance in *35S:MTS1* tomato plants (Figure 4C).

**Figure 5.**
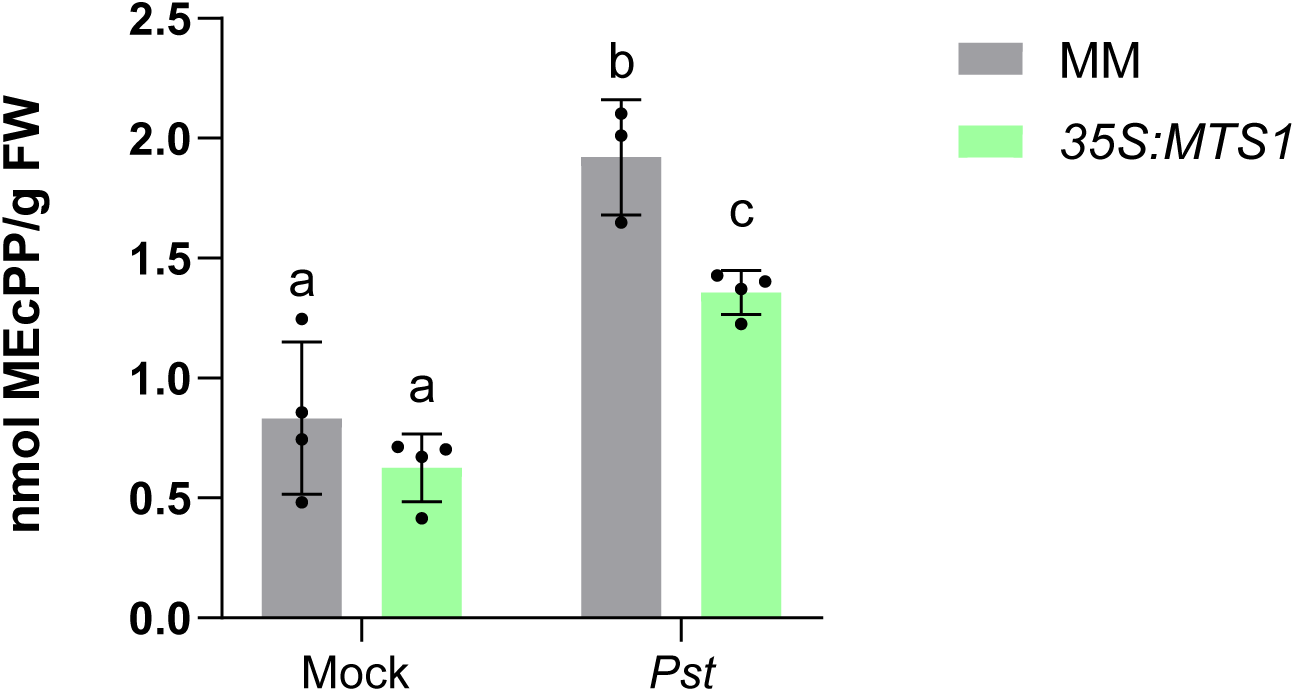
Reduction of MEcPP levels in infected *35S:MTS1* tomato transgenic plants. 2-C-methyl-D-erythritol-2,4-cyclopyrophosphate (MEcPP) content was measured in overexpressing *35S:MTS1* plants and their control MoneyMaker transgenic plants carrying an empty vector (MM) upon infection with *Pst*. Mock represents the non-inoculated plants. Statistically significant differences (*p* < 0.05) between genotypes and infected or mock-treated plants are represented by different letters.

**Figure 6.**
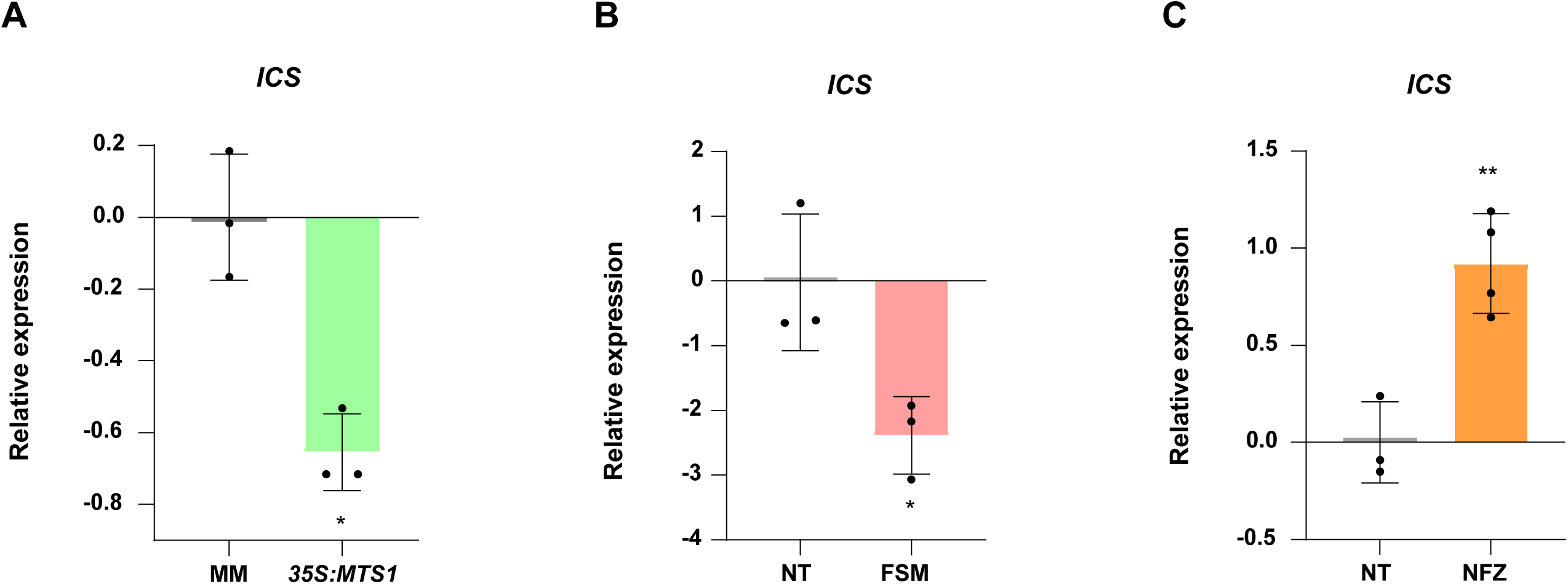
Changes of the relative expression levels of the tomato *ICS* gene in plants with alterations in the MEP-pathway upon bacterial infection in A) *35S:MTS1* plants and their control transgenic plants with empty vector (MM), and in wild type MM plants non-treated (NT) or after **B)** fosmidomycin (FSM) or **C)** norfluorazon (NFZ). Values were normalized to *Actin* gene. Expression levels are represented as mean ± SD of three biological replicates of one representative experiment. Statistically significant differences (*p* < 0.05 or *p* < 0.01) between treated and non-treated are represented by asterisks (*) or (**), respectively.

To better study this interaction between HMTPs and SA biosynthesis, a pharmacological approach was followed to alter the levels of MEcPP, a day before *Pst* infection. Specifically, we used fosmidomycin (FSM) to block the early steps of the MEP pathway and reduce MEcPP production, and norflurazon (NF) to block the consumption of MEP-derived products for carotenoid synthesis and hence predictably cause an increased MEcPP accumulation. FSM specifically inhibits the MEP pathway enzyme deoxyxylulose 5-phosphate reductoisomerase (DXR) whereas NF inhibits the carotenoid pathway enzyme phytoene desaturase (PDS) (Graphical Abstract).

Firstly, we studied the FSM-inhibition effect of the MEP pathway on the *ICS* expression levels in *Pst*-infected tomato plants by RT-qPCR (Figure 6B). Indeed, FSM application caused a statistical decrease in the *ICS* expression levels upon *Pst* infection, thus indicating that the inhibition of the DXR enzyme blocked the MEcPP production and, therefore the transcriptional activation of *ICS*. On the contrary, the inhibition of the carotenoid biosynthesis by NFZ produced a statistically significant higher activation of *ICS* upon *Pst* infection when compared with non-treated infected plants (Figure 6C).

To confirm that regulation of *ICS* produces alterations in SA biosynthesis, we analysed the levels of this phytohormone as well as its hydroxylated (gentisic acid; GA) and methylated forms (methyl-salicylate, MeSA), in both *Pst*-infected transgenic plants with altered *MTS1* levels. Either GA or MeSA are involved in compatible interactions (Belles et al., 2006) and in the activation of SAR response (Bartsch et al., 2010; Lowe-Power et al., 2016), respectively. As shown in Figures 7A and 7B, the levels of SA and GA were statistically lower in *Pst*-infected *35S:MTS1* tomato leaves when compared to its genetic background MM. A significant reduction in GA levels was also measured in mock conditions in these transgenic plants. In contrast, a significant higher SA and GA accumulation was measured in *Pst*-infected *RNAi_MTS1* plants when compared with the corresponding parental plants (Figure 7C and 7D). Thus, the lower *ICS* induction observed in *MTS1* overexpressing lines (Figure 6A) perfectly correlated with the lower levels of SA and GA (Figures 7A and 7B), and with the lower levels of *PR1* upon bacterial infection (Figure 4C). In a similar manner, an opposite MeSA emission was analysed in *35S:MTS1* and *RNAi_MTS1* tomato plants. While *35S:MTS1* tomato leaves showed a lower emission of volatile MeSA after infection, *RNAi_MTS1* plants exhibited a higher amount of MeSA compared to corresponding wild type (Supplemental Figure 1A and C). Moreover, the expression levels of *S5H*, involved in the conversion of SA to GA (Payá et al., 2022), paired with the accumulation levels of the two phenolics in both infected transgenic plants. Thereby, *35S:MTS1* tomato leaves showed a statistical decrease in *S5H* expression levels while *RNAi_MTS1* displayed a slightly significant *S5H* induction after bacterial infection comparing to the parentals (Supplemental Figure 1B and D).

**Figure 7.**
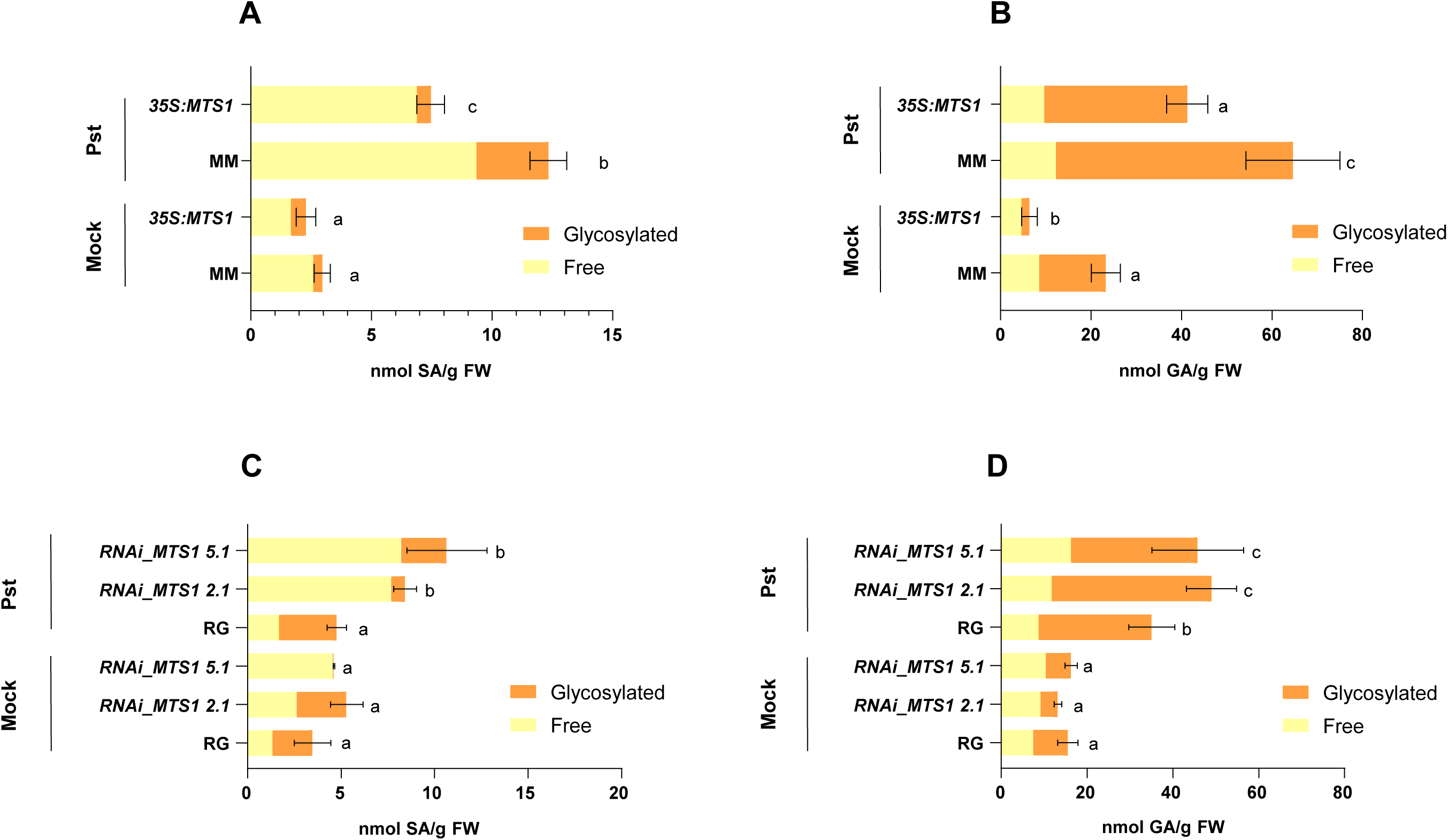
Metabolic cross-talk between monoterpenoids and SA biosynthesis. Levels of free and glycosylated salicylic (SA; left panels) and gentisic acid (GA; right panels) were analyzed in both transgenic tomato plants 24 hours after bacterial (*Pst*) infection. **Upper panels A)** and **B)** show the phenolic content in overexpressing *35S:MTS1* tomato plants and their control plants with the empty vector (MM), and **lower panels C)** and **D)** phenolic content in silencing lines of *RNAi_MTS1* tomato plants and their parental RG. Mock represents the non-inoculated plants. The extracts were analyzed by fluorescence-HPLC. Bars represent the mean ± SD of total levels of a representative experiment (n=5). Significant differences between genotypes and infected or mock-inoculated plants are represented by different letters (*p* < 0.05).

Furthermore, levels of SA and GA were also measured in tomato plants pre-treated either with FSM or with NFZ and infected with *Pst*, which displayed a lower or higher activation of *ICS*, respectively (Figures 6B and 6C). Also consistently, whilst FSM-treated and infected tomato plants accumulated lower levels of SA (Figure 8A) and GA (Figure 8B), NFZ-treated and infected tomato plants accumulated higher levels of both phenolics (Figures 8C and 8D), when compared with non-treated control plants infected with *Pst*.

**Figure 8.**
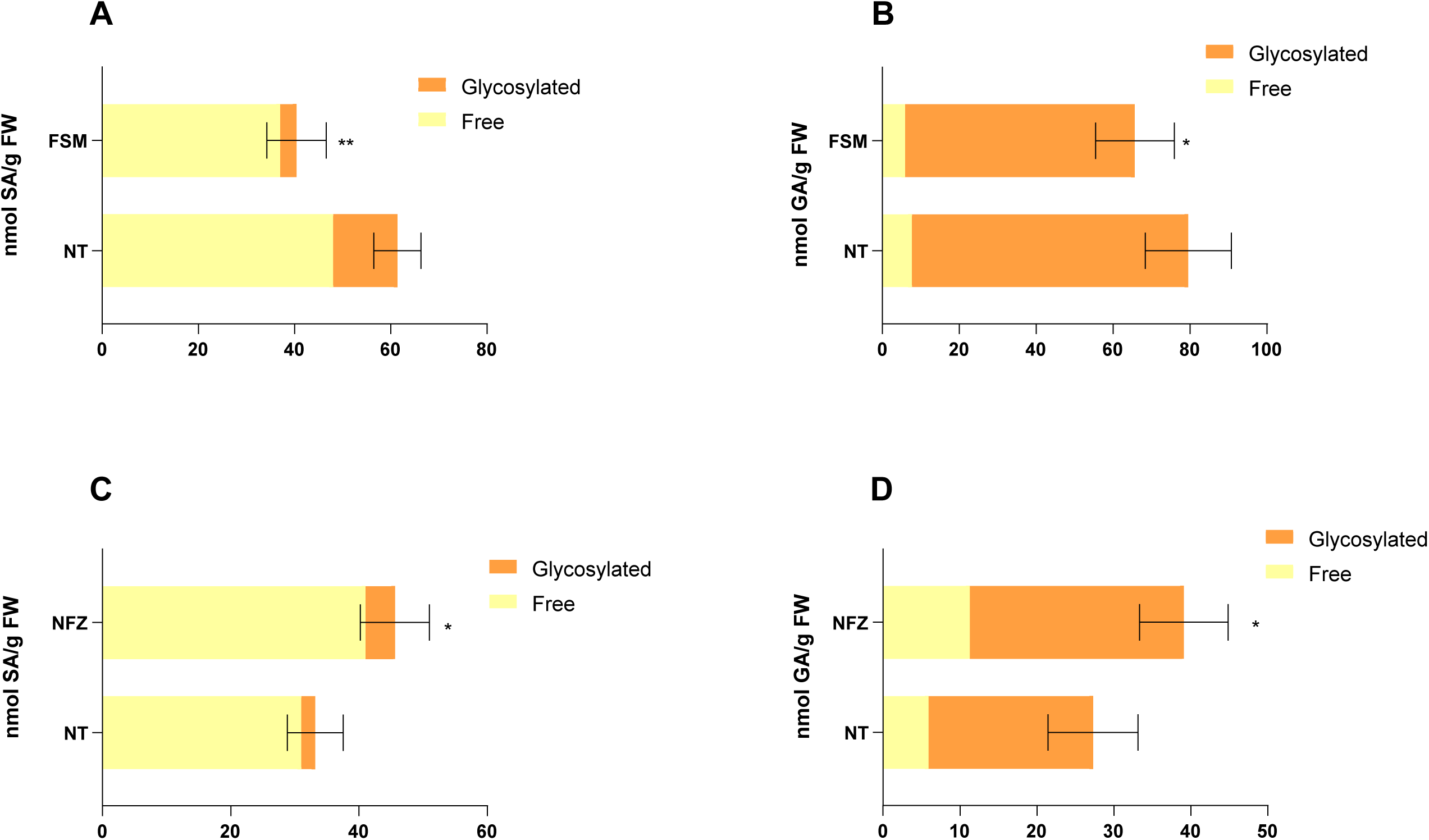
Pharmacological validation of the cross-talk between monoterpenoids an salicylic acid biosynthesis. Free and glycosylated salicylic acid (SA; left panels) and gentisic acid (GA; right panels) levels in different pre-treated MoneyMaker tomato plants displaying alterations in MEP-pathway at 24 hours after bacterial infection. **Upper panels A)** and **B)** show the phenolic content after fosmidomycin (FSM) pre-treatment plants and **lower panels C)** and **D)** the phenolic content after norfluorazon (NFZ) pre-treatment. Non-treated plants (NT). The extracts were analyzed by fluorescence-HPLC. Bars represent the mean ± SD of total levels of a representative experiment (n=5). Statistically significant differences (*p* < 0.05 or *p* < 0.01) between treated and non-treated are represented by asterisks (*) or (**), respectively of total levels of a representative experiment (n=5).

Both genetic and pharmacological approaches appear to indicate that there is a competition for MEcPP by the MEP pathway, which leads to the production of HMTPs, and the *ICS* transcriptional activation, which induces the SA biosynthesis, thus revealing a metabolic crosstalk between both pathways. These findings could explain the fact that, despite the observed statistical differences in HMTPS emission and stomatal aperture ratios, none of the transgenics plants showed a phenotype of resistance (*35S:MTS1*) or susceptibility (*RNAi_MTS1*), since SA levels were inversely affected in these plants, thus confirming the existence of a negative crosstalk between HMTPs and SA during the bacterial infection.

### HMTPs and SA Balance in the Defensive Response of Tomato Plants Against *Pst*

To verify the relevance of the MEP pathway and its connection with SA biosynthesis in plant resistance, FSM-pre-treated or NF-pre-treated tomato plants were infected with *Pst* and the bacterial growth was evaluated. As shown Figure 9A, the number of colonies was statistically higher in FSM-treated plants with respect to the untreated. This result indicates that FSM treatments, which repressed *ICS* (Figure 6B) and reduced SA accumulation (Figure 8A), increase the susceptibility of tomato plants against *Pst*. On the other hand, NF treatments produced enhanced resistance (Figure 9B), due to the activation of *ICS* (Figure 6C) and SA accumulation (Figure 8C).

**Figure 9.**
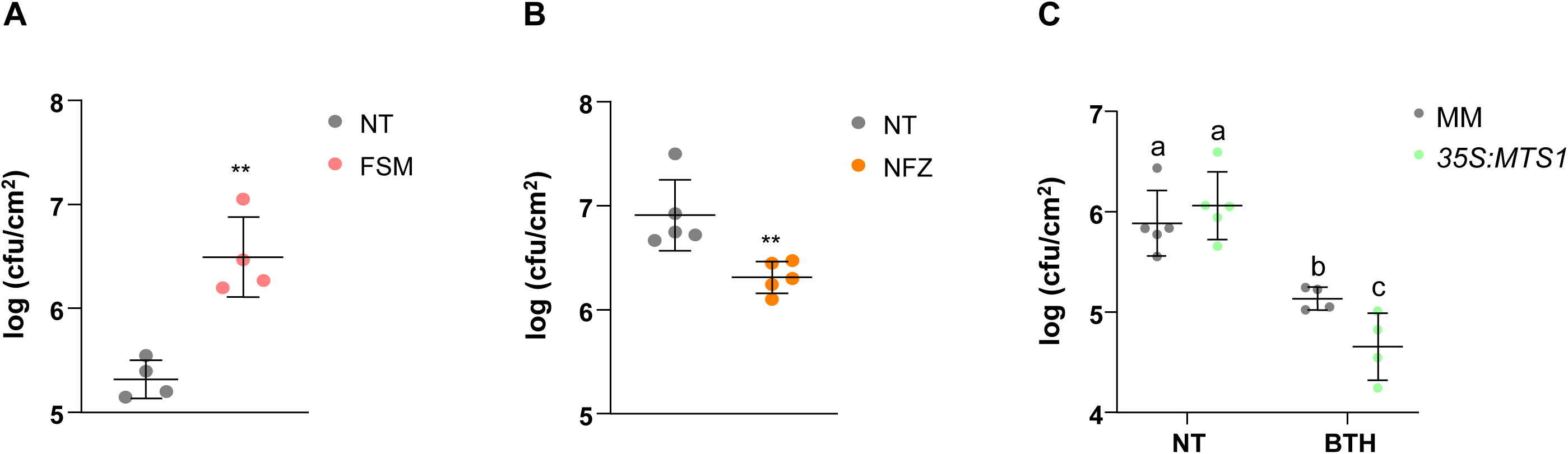
Role of the MEP pathway on the tomato resistance to bacteria. **A)** and **B)** Bacterial content in MoneyMaker tomato plants pre-treated either with fosmidomycin (FSM) or norfluorazon (NFZ), respectively. Tomato plants were inoculated with *Pst* by immersion and leaf samples were taken 24 hours after bacterial infection. Data are presented as means ± SD of a representative experiment (n=5). Statistically significant differences (*p* < 0.01) between treated and non-treated are represented by asterisks (**) **C)** Effect of benzothiadiazole (BTH) treatments in the resistance of *35S:MTS1* transgenic plants and their corresponding control plants with the empty vector (MM). Data are presented as means ± SD of a representative experiment (n=5). Statistically significant differences (*p* < 0.05) between genotypes and BTH or non-treated (NT) plants are represented by different letters.

Both treatments confirm the relation between the MEP pathway and SA biosynthesis and highlight the importance of HMTPs and SA in the plant resistance. These studies clarify the importance of the MEP pathway in the tomato resistance through MEcPP, a compound participating in both the transcriptional stimulation of *ICS* and the biosynthesis of HMTPs.

Finally, to confirm the role of HMTPs in plant defence against bacteria, treatments with the SA functional analogue benzothiadiazole (BTH), used as chemical activator of plant resistance by activating SAR (Lawton et al., 1996), were carried out in the transgenic *35S:MTS1* tomato plants. Applications of 1 mM BTH in transgenic *35S:MTS1* plants produced a significant resistance when compared with corresponding treated parental (Figure 9C). This resistance was accompanied by a restored accumulation of *PR1* in the *35S:MTS1* plants (Supplemental Figure 2) when compared with those previously obtained (Figure 4C).

### HMTPs participate in the communication between tomato plants

The role of VOCs, in intra- and inter-communication between plants is well-known (Baldwin I et al., 2002; Zimmermann et al., 2009). As long-distance signals, VOCs can trigger systemic stress responses in distant plants. As Figure 10 shows, when tomato plants (receivers) were cohabited with *35S:MTS1* plants which over-emit HMTPs (emitters; Figure 3C), stomatal closure was observed in those receiver plants. Moreover, the observed effect occurred in a dose-dependent manner, since the stomata closure was less pronounced when two emitter plants were used (Figure 10A) in comparison with the results obtained with four emitter plants (Figure 10B). Besides, *35S:MTS1* emitter plants activated the plant defence response in the receiver plants, as levels of *PR1* showed (Figure 10C). Therefore, our results indicate that HMTPs may play an important role not only in the intra-but also in the inter-plant immune signalling.

**Figure 10.**
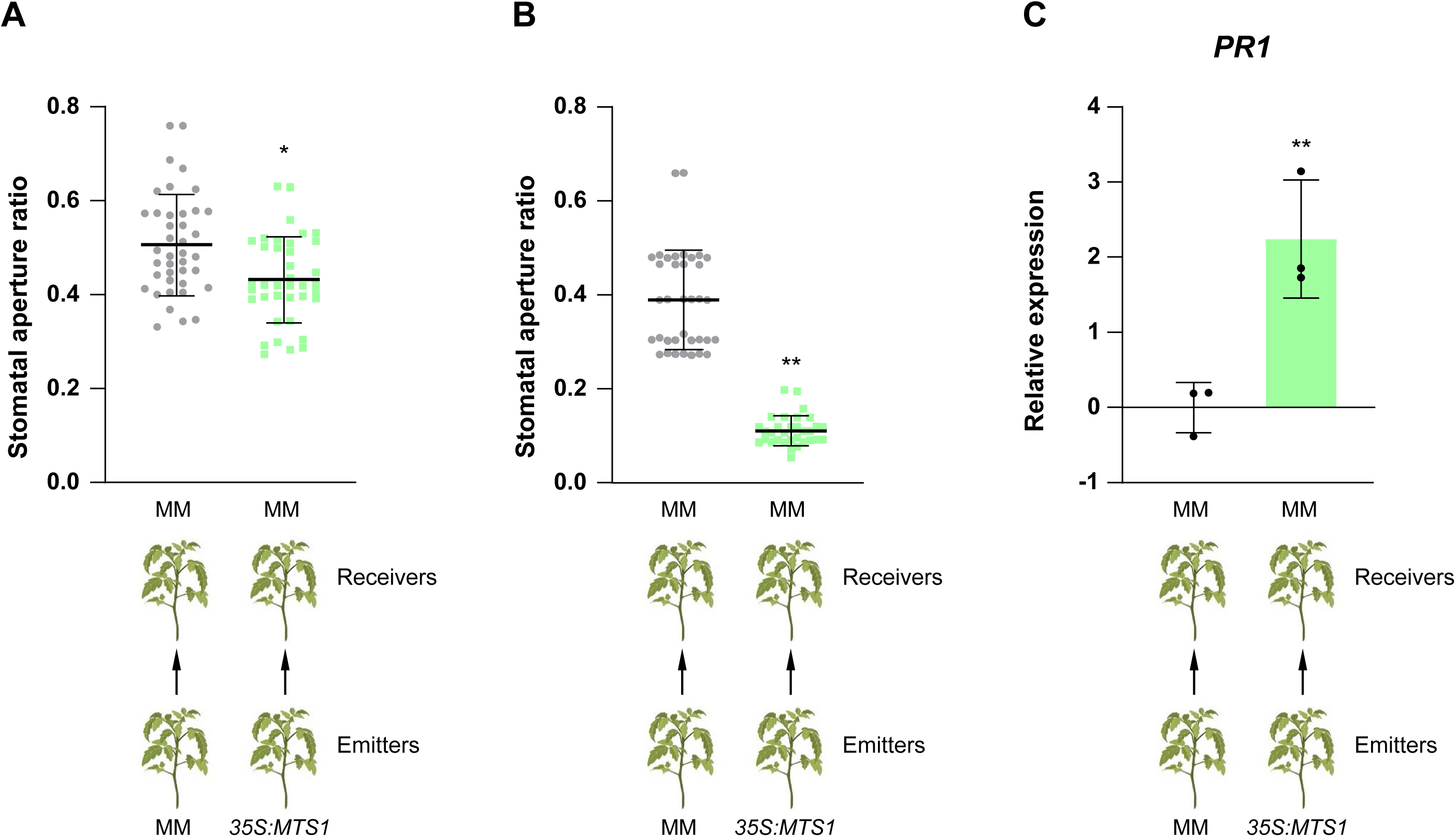
Interplant communication using *35S:MTS1* plants as emitters. Tomato MoneyMaker plants (MM; “Receivers”) were placed in closed chambers in the presence of *MTS1* overexpressing plants (“Emitters”) or their corresponding control plants with the empty vector MM as control Emitters, and stomatal aperture ratio of three biological replicates was measured in receivers of tomato plants after cohabitation for 24 h. Two emitters *vs.* 2 receivers were used in **(A)**, and four emitters *vs.* 2 receivers were used in **(B)**. The relative expression of tomato *PR1* gene **(C)** was analyzed by qRT-PCR in MM receiver plants after cohabitation either with *35S:MTS1* or MM emitters. Two emitters *vs.* 2 receivers were used. Values were normalized to Actin gene. Expression levels are represented as mean ± SD of three biological replicates of one representative experiment. Statistically significant differences (*p* < 0.05 or *p* < 0.01) between treated and non-treated plants are represented by asterisks (*) or double asterisks (**), respectively.

## DISCUSSION

Terpenoids constitute a highly diverse class of chemical compounds, which are abundantly produced across the Plant Kingdom (Zhou and Pichersky; 2020). Particularly, monoterpenes are implicated in the plant defence response, being their defensive role classically associated with plant-herbivore interaction. However, its role in plant defence against pathogens is earning interest (Vlot et al., 2021). In this sense, some HMTPs as α-terpineol, 4-terpineol or linalool have been described as differentially emitted VOCs during ETI establishment triggered by *Pst* in tomato plants (López-Gresa et al.,2017). Here we study the defensive role of these HMTPs and its convergence with the SA-mediated immunity in tomato plants.

Stomata participate in the gas exchange that allows transpiration and photosynthesis, being indirectly involved in plant immunity as these apertures act as entry route for pathogens to the vegetal tissue (Melotto et al., 2009). Besides, the activation of plant defence leads to the induction of the so-called Pathogenesis Related (PR) proteins, being PR1 the conventional SA-related marker (Saijo and Loo, 2020). We have observed that all the analyzed HMTPs treatments provoked both stomatal closure and *PR1* induction, unlike the non-hydroxylated limonene, thus pointing out the importance of the hydroxylation for the activation of the plant defence response (Figure 1). To the best of our knowledge, this is the first time that HMTPs have been reported as plant stomata closers. Only, previous investigations of Rai *et al*. 2003 evidence that volatile essential oils from *Pinsepia utilis* inhibit stomatal opening in *Vicia faba*. Particularly, α-terpineol attracts a great interest as an antioxidant, anticancer, anticonvulsant, antiulcer, antihypertensive, or anti-nociceptive compound (Khaleel et al., 2018), being the most efficient volatile in terms of stomatal closure (Figure 1A). To elucidate the defensive role of HMTPs, the capacity to activate plant resistance and the stomata closure process triggered by α-terpineol treatments were evaluated. This monoterpenoid induced *PR1* expression (Figure 1B) and tomato resistance against *Pst* (Figure 1C), confirming its defensive role. Our results are in accordance with those previously described in *Arabidopsis thaliana,* where a mixture of monoterpenes α-pinene and β-pinene induced resistance (Riedlmeier et al., 2017), although their capacity to induce stomatal closure was not explored. In other studies, monoterpenes had already been associated with the defensive response such as geraniol and its potential antibacterial effects against *Xanthomonas oryzae* pv. *oryzae*, which causes rice bacterial blight (Kiyama et al., 2021).

The effective stomata closure after α-terpineol treatment was confirmed by the lower loss of water observed in treated plants (Figure 2A), being the α-terpineol active at concentrations similar to those used for ABA (Figure 2B), a plant hormone with a central role in the regulation of stomatal movements under water-deficit conditions (Hsu et al, 2021). Surprisingly, the observed stomata closure occurred in a SA and ABA-independent manner (Figures 2C and 2D), both positive regulators of the stomatal immunity (Melotto et al., 2006, 2017; Su et al., 2017). Similar results were previously described for oxylipins, which participate in the stomatal immunity in an ABA-independent process (Montillet et al, 2013), or for the (*Z*)-3-hexenyl butyrate, a volatile compound also emitted by tomato plants displaying ETI which has been described to close stomata in a SA and ABA-independent manner (López-Gresa et al., 2018; Payá et al., under review). Our results reinforce the existence of ABA independent pathways in stomatal immunity.

After the pharmacological approaches, the capacity of HMTPs to induce resistance was analyzed in transgenic plants with altered levels of *MTS1* expression. For this purpose, we used the previously described *35S:MTS1* plants (Van Schie et al., 2007) which over-emit HMTPs, and we generated *RNAi_MTS1* plants with reduce emission levels of HMTPs (Figure 3). Surprisingly, levels of emission of HMTPs did not correlate with plant resistance Figures 4B and 4E, even though the capacity of the HMTPs to regulate stomata was maintained in these transgenic plants (Figures 4A and 4D). Therefore, our results appeared to indicate the ability of HMTPs to confer resistance was not only due to stomatal closure. For a further characterization, *PR1* expression levels were measured, observing a reverse correlation between HMTPs emission and the expression of this defence marker gene (Figures 4C and 4F). These results suggested that, somehow, HMPTs alter *PR1* expression and, therefore, the SA-mediated response.

Monoterpenes and both SA biosynthesis and signalling have been previously related in Arabidopsis. Particularly, pinene-induced resistance was described to be dependent of SA biosynthesis and signalling (Riedlmeier et al., 2017). Besides, *CSB3* which encodes a 1-hydroxy-2-methyl-2-butenyl 4-diphosphate synthase participating in the MEP pathway, is expressed constitutively in healthy plants, and shows repression in response to bacterial infection, being described as a point of metabolic convergence between MEP and SA-mediated disease resistance to biotrophic pathogens (Gil et al., 2005). Finally, the MEP pathway is connected to SA through *ICS* expression by the retrograde signaller MEcPP. This compound is a precursor of isoprenoids which elicits the expression of selected stress-responsive nuclear-encoded plastidial proteins as *ICS*, the main producer of SA (Xiao et al., 2012).

Accordingly, we observed that *35S:MTS1* transgenic plants displayed lower levels of MEcPP accumulation (Figure 5), a reduced induction of *ICS* (Figure 6A), lower levels of SA (Figure 7A) and therefore lowered PR1 activation (Figure 4B) upon *Pst* infection. Conversely, higher levels of SA (Figure 7C) and enhanced *PR1* activation (Figure 4E) were detected in *RNAi_MTS1* plants. These findings could explain our observed phenotypes in tomato *MTS1* transgenic plants, since neither a resistance nor a susceptibility were obtained (Figures 4C and 4D), probably due to the alteration of the MEP pathway. In the case of *35S:MTS1* plants, the constitutive promoter would cause a depletion of MEP pathway precursors which are routed into the biosynthesis of monoterpenes. Therefore, a lower amount of MEcPP could act as retrograde signal, reducing the *ICS* transcriptional activation. On the contrary, in *RNAi_MTS1* plants an accumulation of the MEP precursors, including MEcPP, occurs and *ICS* is then induced. The absence of phenotype in both cases account for the important role of both SA and HMTPs in tomato defence against bacteria.

To further confirm this last hypothesis, we performed two pharmacological approaches. Firstly, application of FSM, a molecule that targets the second enzyme of the MEP pathway reducing the levels of the final products (Laule et al., 2003; Graphical Abstract), was used to test the impact of eliminating both monoterpenoids and the transcriptional regulation of *ICS* in the defensive response. After FSM treatment, infected tomato plants displayed a down-regulation of *ICS* (Figure 6B), lower levels of SA (Figure 8A) and consequently a higher susceptibility (Figures 9A), confirming the connection between the MEP pathway and SA-mediated defence. Furthermore, a NFZ treatment, which blocks the carotenoid biosynthesis (Di et al., 2023; Graphical Abstract), was performed to check if an accumulation of MEP precursors could modulate SA regulation. Contrary to FSM treatment, upregulation of *ICS* (Figure 6C), higher SA accumulation (Figure 8C) and resistance to bacteria were observed (Figures 9B), corroborating the aforementioned crosstalk between HMTPs and SA.

In conclusion, MEcPP is revealed as a key metabolite in tomato defence against biotic stress since it is necessary for both monoterpenoid biosynthesis and transcriptional activation of *ICS*. The lower levels of MEcPP detected in *35S:MTS1* confirmed this biosynthetic crosstalk (Figure 5). In addition, the hyper susceptibility and resistance phenotype observed in tomato plants after treatments with fosmidomycin or norfluorazon respectively, as well as enhanced resistance of *35S:MTS1* plants after BTH treatments against *Pst* (Figure 9C) indicate that both HMTPs and SA are required for resistance induction in tomato plants. Furthermore, the biosynthetic regulation of both pathways mediated by MEcPP is key in the context of the defensive response and must be fine-tuned to optimize defence and fitness.

Finally, we have observed that *35S:MTS1* tomato plants induce stomata closure and *PR1* expression in neighbour receiving tomato plants, thus indicating that HMTPs play an important role in inter-plant communication (Figure 10). Similar results were observed in Arabidopsis, since monoterpenes contributed to defence related plant-to-plant communication (Riedlmeier et al., 2017). MeSA and nonanal are VOCs which have also been described to trigger plant defence (Shulaev et al., 1997; Yi et al., 2009). However, since *35S:MTS1* tomato plants emitted lower levels of MeSA (Supplemental Figure 1A), the induction of plant defence in receiving plants is probably due to the HMTPs, reinforcing their role in the defence related plant communication.

To sum up, by using pharmacological and genetical approaches we have demonstrated that HMTPs play an important defensive role in tomato, contributing to stomatal closure and the activation of the defence response not only within the plant, but also in the neighbouring plants, therefore acting as signal molecules for intra- and inter-plant communication. Moreover, our results have revealed a metabolic crosstalk between MEP and SA pathways, occurring through MEcPP competition (Graphical Abstract).

## MATERIALS AND METHODS

### Vector construction and tomato transformation

In order to generate the *MTS1*-silenced transgenic tomato plants, the method described by Helliwell and Waterhouse was followed (Helliwell and Waterhouse, 2003). Briefly, a selected 407 bp sequence of MTS1 was amplified from the full-length cDNA clone using the forward primer 5′-GGCTCGAGTCTAGAATGGTTTCAATATTGAGTAAC -3′, which introduced restriction sites *Xho*I and *Xba*I, and the reverse primer 5′-CCGAATTCGGATCCCTCCTCATAATTTGCATAATTTCATC-3′, which added restriction sites *Bam*HI and *Eco*RI. The PCR product was first cloned in the pGEM-T Easy vector (Promega) and sequenced. After digestion with the appropriate restriction enzymes and purification, the two *MTS1* fragments were subcloned into the pHANNIBAL vector in both the sense and antisense orientations. Finally, the constructs in pHANNIBAL were subcloned as a *Not*I flanked fragment into a binary vector pART27 to produce highly effective intron-containing “hairpin” RNA silencing constructs. This vector carries the neomycin phosphotransferase gene (*NPT II*) as a transgenic selectable marker.

The transformed LBA4404 *Agrobacterium tumefaciens* carrying pART27-MTS1 was co-cultured with tomato RG cotyledons to generate the RNAi MTS1-silenced transgenic tomato plants (*RNAi_MTS1*). The explant preparation, selection and regeneration methods followed those published by Ellul and co-workers (Ellul et al., 2003). The tomato transformants were selected in kanamycin-containing medium and propagated in soil. RG tomato wild-type plants regenerated *in vitro* from cotyledons under the same conditions as the transgenic lines were used as controls in subsequent analyses. The transgenic plants generated in this study have been identified and characterised in our laboratory and are to be used exclusively for research purposes.

### Plant material and growth conditions

For the purposes of this study, we used different tomato genotypes: (i) *NahG* (Brading et al., 2000) and its parental Moneymaker (MM, kindly provided by Prof. Jonathan Jones, The Sainsbury Laboratory, Norwich, UK); (ii) *35S:MTS1* (Van Schie et al., 2007) and the parental with the empty vector pGreen (gently provided by Prof. Schuurink Swammerdam Institute for Life Sciences, Department of Plant Physiology, University of Amsterdam, The Netherlands); (iii) *flacca* mutants and its parental Lukulus (all of them kindly provided by Dr. Jorge Lozano, Instituto de Biología Molecular y Celular de Plantas UPV-CSIC, Valencia, Spain); (iv) and *RNAi_MTS1* plants generated in our laboratory and its parental RG which contain the *Pto* resistance gene (a gift from Dr. Selena Giménez, Centro Nacional de Biotecnología, Madrid, Spain). A mixture of sodium hypochlorite:distilled H_2_O (1:1) was used for the sterilization and sequential washings of 5, 10, and 15 min for the total removal of hypochlorite. Seeds germinated were placed in 12 cm-diameter pots with vermiculite and peat. The greenhouse conditions were the following ones: a relative humidity of 50 approximately and a 16/8 h (26 ◦C) light/dark photoperiod.

### HMTPs treatments and interplant communication

Treatments were carried out in 4-week-old tomato plants. Tomato plants were put into 121 L methacrylate chambers containing hydrophilic cotton buds with 5 μM monoterpenoid with 0.05 % (v/v) Tween-20 or distilled water. Methacrylate chambers were hermetically sealed during 24 h. For spray treatments, tomato plants were pre-treated by spray with 2 mM monoterpenoid with 0.05 % (v/v) Tween-20 or distilled water.

For interplant communication assays, 2 or 4 emitter plants were cohabitated with 2 receiver plants in the mentioned methacrylate chambers for 24 h. Plant material was only collected from the MM wild type receiver plants.

### Inhibitors and BTH treatments

28-days-old MM plants were sprayed either with 50 µM fosmidomycin with 0.05 % (v/v) Silwet detergent solution or with 100 µM norflurazon with 0.05 % (v/v) Silwet detergent solution. 1Mm BTH was applied 24 h before the infection with 0.05 % (v/v) Silwet detergent solution.

### Bacteria inoculation and CFU determination

The bacterial strain used in this study was *Pseudomonas syringæ pv. tomato* DC3000 with deletions in genes *avrPto* and *avrPtoB* (*Pst*) (Lin and Martin, 2005; Ntoukakis et al., 2009). Bacterial growth conditions and plant inoculation were performed as previously described (López-Gresa et al., 2018).

Briefly, for colony formation units (CFU) measurements three leaf disks (1 cm^2^ each) were grounded and serial dilutions of the infected tissue were cultured on King’s B agar medium containing rifampicin. CFU were counted after 48 h at 28 °C.

### Stomatal aperture

Stomatal samples for the observation of aperture ratio were collected with a layer of nail polish in the abaxial part of the leaves and the epidermis peels were placed on slides for their observation with a Leica DC5000 microscope (Leica Microsystems S.L.U.). In total, 50 stomata of each condition were analyzed using the NIH’s ImageJ software. Thus, several photos were taken from different regions of the tomato leaves. Stomatal aperture ratio was calculated as width/length.

### Weight loss experiments

Around 15-25 tomato plants were grown in *in vitro* conditions treated with 5 µM α-terpineol for 24 h. The aerial and root parts were weighted every minute for 3 h at room temperature. After 24 hours, the dry weight was measured.

### RNA isolation and quantitative RT-PCR analysis

The RNA extraction and conversion to cDNA of tomato leaves was carried out using columnkit based on silica membranes (Macherey-Nagel XXX GmbH, Germany) following the manufacturer’s protocol. cDNA from a microgram of RNA was obtained using a PrimeScript RT reagent kit (Perfect Real Time, Takara Bio Inc., Otsu, Shiga, Japan) following its instructions. Quantitative real-time PCRs were performed as previously described (Campos et al., 2014). In each plate of a 96-wells plates a reaction took place in a final volume of 10 µL. SYBR R Green PCR Master Mix (Applied Biosystems) was used as the fluorescence marker and actin gene as the endogenous reference gene. The PCR primers were designed using the online service Primer3 (http://primer3.ut.ee/) and are listed in Table S1.

### SA and GA measurements

For the extraction of SA and GA, 0.5 g of frozen homogenized leaves were resuspended in methanol which contained 25 mM *o*-anisic acid as internal standard. The supernatant, after a 10 min-centrifugation and 10 min-ultrasounds, was divided into two eppendorfs and dried using a nitrogen flow. For the analysis of total and glycosylated SA and GA, the protocol described by Vazquez Prol F *et al*. (Vázquez Prol et al.,2021) was carried out. Quantification of SA and GA was obtained using a calibration curve based on the internal standard.

### GC-MS

For the analysis of volatile organic compounds, a mix of 1 mL of CaCl_2_ 6 M and 100 µL of 750 mM EDTA at a pH of 7.5 was added to 100 mg of pulverize tomato leaves in 10 mL glass vial. The vials were air-tight sealed and sonicated for 5 min. Volatile compounds extraction was performed by Head Space Solid-Phase Microextraction (HS-SPME) (López-Gresa et al., 2017). Enhanced ChemStation software (Agilent) was the program used to obtain and analyze the chromatograms and mass spectra which has its own database to compare the different ion and retention times with pure compounds. Quantification of monoterpenes was performed elaborating a method in Agilent using the most abundant ion and retention time and calculating the area on the chromatogram.

### MEcPP measurements

MEcPP was quantified according to the protocol described in Baidoo et al., 2014 (Baidoo et al., 2014) with minor modifications.100 mg of frozen homogeneized tissue were extracted in 13 mM ammonium acetate buffer pH 5.5. The extract was dried under a nitrogen steam and resuspended in the UPLC mobile phase (73% acetonitrile, 27% 50 mM ammonium carbonate in water)

MEcPP quantification was performed using a Orbitrap Exploris 120 mass spectrometer coupled with a Vanquish UHPLC System (Thermo Fisher Scientific, Waltham, MA, USA). LC was carried out by reverse-phase ultraperformance liquid chromatography using a BEHAmide column (1.7 uM particle size, dimensions 2.1 x 150 mm) (Waters Corp., Mildford, MA, USA)

The mobile phase was 73% acetonitrile, 27% 50 mM ammonium carbonate in water. Samples were run in isocratic mode for 14 minutes. The flow rate was 0.2 ml/min and the injection volume was 5 uL. The column temperature was set at 30°C.

Ionisation was performed with heated electrospray ionization (H-ESI) in positive mode. Samples were acquired in full scan mode (resolution set at 120000 measured at FWHM). Methionine sulfone and D4-succinic acid were used as internal standards. For absolute quantification a standard curve was performed with MEcPP chemical standard. Data processing was performed with TraceFinder software (Thermo Scientific, Waltham, MA, USA).

### Western Blot

Protein extracts for immunodetection experiments were prepared from MM plants treated and non-treated with α-terpineol for 24h. Material (100 mg) for direct western-blot analysis was extracted in 23 Laemmli buffer (125 mM TrisHCl, Ph 6.8,4%[w/v] SDS, 20% [v/v] glycerol, 2% [v/v] mercaptoethanol, and 0.001% [w/v] bromophenol blue), and proteins were run on a 10% SDS-PAGE gel and analyzed by immunoblotting. Proteins were transferred onto Immobilon-P membranes (Millipore) and probed with anti-rabbit-peroxidase (Jacksons). Immunodetection of defensive protein PR1 was performed using antiPR1 antibody. Antibodies were used at a 1:20000 dilution. Detection was performed using the ECL Advance Western Blotting Chemiluminiscent Detection Kit (GE Healthcare). Image capture was done using the image analyzer LAS3000, and quantification of the protein signal was done using Image Guache V4.0 software.

## ACKNOWLEDGMENTS

We would like to thank IBMCP Metabolomics Platform (Valencia, Spain), especially to Teresa Caballero for her excellent technical support in the VOCs quantification and Dr. Ana Espinosa for the MEcPP quantification.

This work was supported by Grant PID2020-116765RB-I00 funded by MCIN/ AEI/10.13039/501100011033/ and Grant PROMETEU/2021/056 by Generalitat Valenciana. C.P. was a recipient of a predoctoral contract of the Generalitat Valenciana (ACIF/2019/187) and J.P-P is a recipient of a predoctoral contract of the Ministerio de Universidades e Investigacion (FPU21/00259).

## SUPPLEMENTAL DATA FILES

Supplemental Figure 1

Supplemental Figure 2 Table S1

