## Supplemental Figure 1 for "Metabolic crosstalk between hydroxylated monoterpenes and salicylic acid in tomato defence response against *Pseudomonas syringae* pv *tomato*"

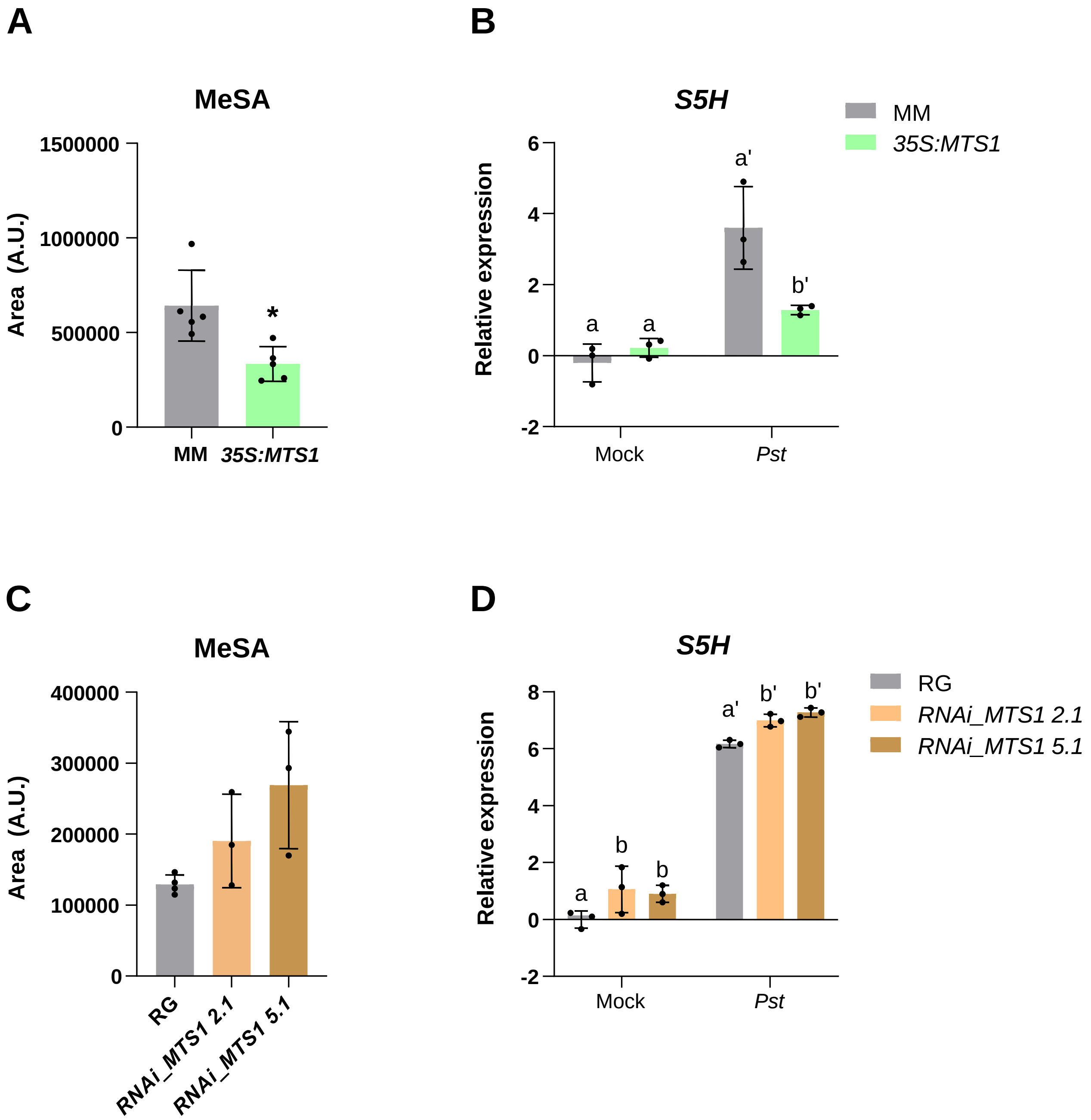


**Supplemental Figure 1. Changes in the methyl salicylate (MeSA) content and in the relative expression levels of the tomato *S5H* gene in plants with alterations in *MTS1* upon bacterial infection. Left panels.** Levels of MeSA in **A)** *MTS1* overexpressing plants and their parentals carrying an empty vector (MM) and in **C)** both silencing lines of *RNAi_MTS1* (2.1 and 5.1) and their parental (RG). Statistically significant differences with non-transgenic plants are represented with asterisk (*) with $p < 0.05$. **Right panels.** The qRT-PCR expression analysis of the tomato *S5H* gene is shown in **B)** for *35S:MTS1* plants and their parental plants (MM), and in **D)** for both lines of *RNAi_MTS1* and their parental (RG). Expression values were normalized to Actin gene. Expression levels are represented as mean ± SD of three biological replicates of one representative experiment. Letters represent statistically significant differences ($p < 0.05$) between genotypes and infected or mock-treated plants.
