## Supplemental Figure 1 for "Metabolic crosstalk between hydroxylated monoterpenes and salicylic acid in tomato defence response against *Pseudomonas syringae* pv *tomato*"

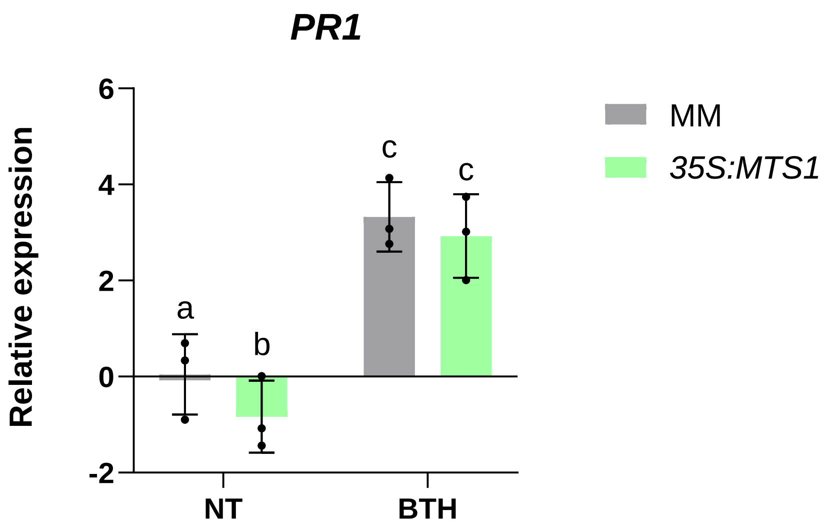


**Supplemental Figure 2. Effect of benzothiadiazole (BTH) treatments in the resistance of *35S:MTS1* transgenic plants.** Relative expression analysis by qRT-PCR of the tomato *PR1* gene in *35S:MTS1* plants and their control transgenic plants with empty vector (MM) after BTH and water (NT; non-treated) treatments. Values were normalized to Actin gene. Expression levels are represented as mean ± SD of three biological replicates of one representative experiment. Statistically significant differences ($p < 0.05$) between genotypes and treated plants are represented by different letters.
