## Supplementary material for "Metabolic crosstalk between hydroxylated monoterpenes and salicylic acid in tomato defence response against *Pseudomonas syringae* pv *tomato*": Table S1

| *Gen* | *Foward Primer* | *Reverse Primer* |
|---|---|---|
| ***ICS*** | **5’ TGC CTC ATG GAC ATA CCA GA 3’** | **5’ TAT GCG AAT GGG GAT TTT TTC 3’** |
| ***PR1*** | **5’ ACT CAA GTA GTC TGG CGC AAC TCA 3’** | **5’ AGT AAG GAC GTT GTC CGA TCG AGT 3’** |
| ***S5H*** | **5’ GGG ATG TCC CGG AAG TAA GT 3’** | **5’ GGC ATT GGA TGG GAT ATT CA 3’** |
| ***MTS1*** | **5’ TGG TGG TCA CCT TCA AGA GA 3’** | **5’ GCC TTG TGG AAA TAG GA 3’** |
| ***Actin*** | **5’ CTA GGC TGG GTT CGC AGG AGA TGA TGC 3’** | **5’ GTC TTT TTG ACC CAT ACC CAC CAT CAC AC 3’** |

**Table S1.** Primer sequences used for qRT-PCR analyses.
